# Ornithine is the central intermediate in the arginine degradative pathway and its regulation in *Bacillus subtilis*

**DOI:** 10.1101/2023.04.21.537655

**Authors:** Robert Warneke, Tim Benedict Garbers, Christina Herzberg, Georg Aschenbrandt, Ralf Ficner, Jörg Stülke

## Abstract

**ABSTRACT:** The Gram-positive model bacterium *Bacillus subtilis* is able to utilize a variety of proteinogenic and non proteinogenic amino acids as sources of carbon, energy and nitrogen. The utilization of the amino acids arginine, citrulline and ornithine is catalyzed by enzymes that are encoded in the *rocABC* and *rocDEF* operons and by the *rocG* gene. Expression of these genes is under control of the alternative sigma factor SigL. RNA polymerase associated to this sigma factor depends on an ATP-hydrolyzing transcription activator to initiate transcription. The RocR protein acts as transcription activator for the *roc* genes. In this work, we have studied the contributions of all enzymes of the Roc pathway to the degradation of arginine, citrulline and ornithine. This identified the previously uncharacterized RocB protein as responsible for the conversion of citrulline to ornithine. *In vitro* assays with the purified enzyme suggest that it acts as a manganese-dependent N-carbamoyl-L-ornithine hydrolase that cleaves citrulline to ornithine and carbamate. So far, the molecular effector that triggers transcription activation by RocR has not been unequivocally identified. Using a combination of transcription reporter assays and biochemical experiments we demonstrate that ornithine is the molecular inducer for RocR activity. Our work suggests that binding of ATP to RocR triggers its hexamerization, and binding of ornithine then allows ATP hydrolysis and activation of *roc* gene transcription. Thus, ornithine is the central molecule of the *roc* degradative pathway as it is the common intermediate of arginine and citrulline degradation and the molecular effector for the transcription regulator RocR.

**IMPORTANCE:** Amino acids serve as building blocks for protein biosynthesis in each living cell but can also be used as sources of carbon, energy and nitrogen. In this work we have identified ornithine as the central player in the utilization of arginine, citrulline and ornithine in the Gram-positive bacterium *B. subtilis*. Ornithine is the common intermediate after the first steps of arginine and citrulline degradation. We have identified the so far uncharacterized protein RocB as the enzyme responsible for the cleavage of citrulline to ornithine and carbamate. Moreover, we demonstrate that ornithine is the molecular effector that triggers ATPase activity of the transcription factor RocR. Binding of ornithine to RocR and the subsequent hydrolysis of ATP allow a functional interaction with the alternative sigma factor SigL and subsequent transcription activation of all genes of the degradative pathway.

## INTRODUCTION

Amino acids are the essential building blocks for the biosynthesis of proteins and many other cellular components. Moreover, many bacteria can use amino acids as a nutrient, and are able to utilize them as a single source of carbon, nitrogen, and energy. Cells can acquire amino acids as a nutrient by uptake from the environment or by the degradation of external peptides or proteins. The Gram-positive model bacterium *Bacillus subtilis* is capable of utilizing a wide range of amino acids, among them glutamate, glutamine, proline, histidine, asparagine and aspartate, serine, alanine, threonine, the branched-chain amino acids, and arginine. The *B. subtilis* genome encodes all the functions for the uptake and utilization of these amino acids which are degraded to intermediates of central carbon metabolism such as pyruvate, 2-oxoglutarate, or oxaloacetate.

Among the amino acids used by *B. subtilis* are arginine and the metabolically closely related non proteinogenic amino acids ornithine and citrulline (1). In addition to glutamate, the general amino group donor of the cell, and 2-oxoglutarate which feeds into the citric acid cycle, the degradation of arginine also produces reducing power in the form of NADH + H^+^ which can be used for ATP synthesis in respiration (see Fig. 1).

**Fig. 1.**
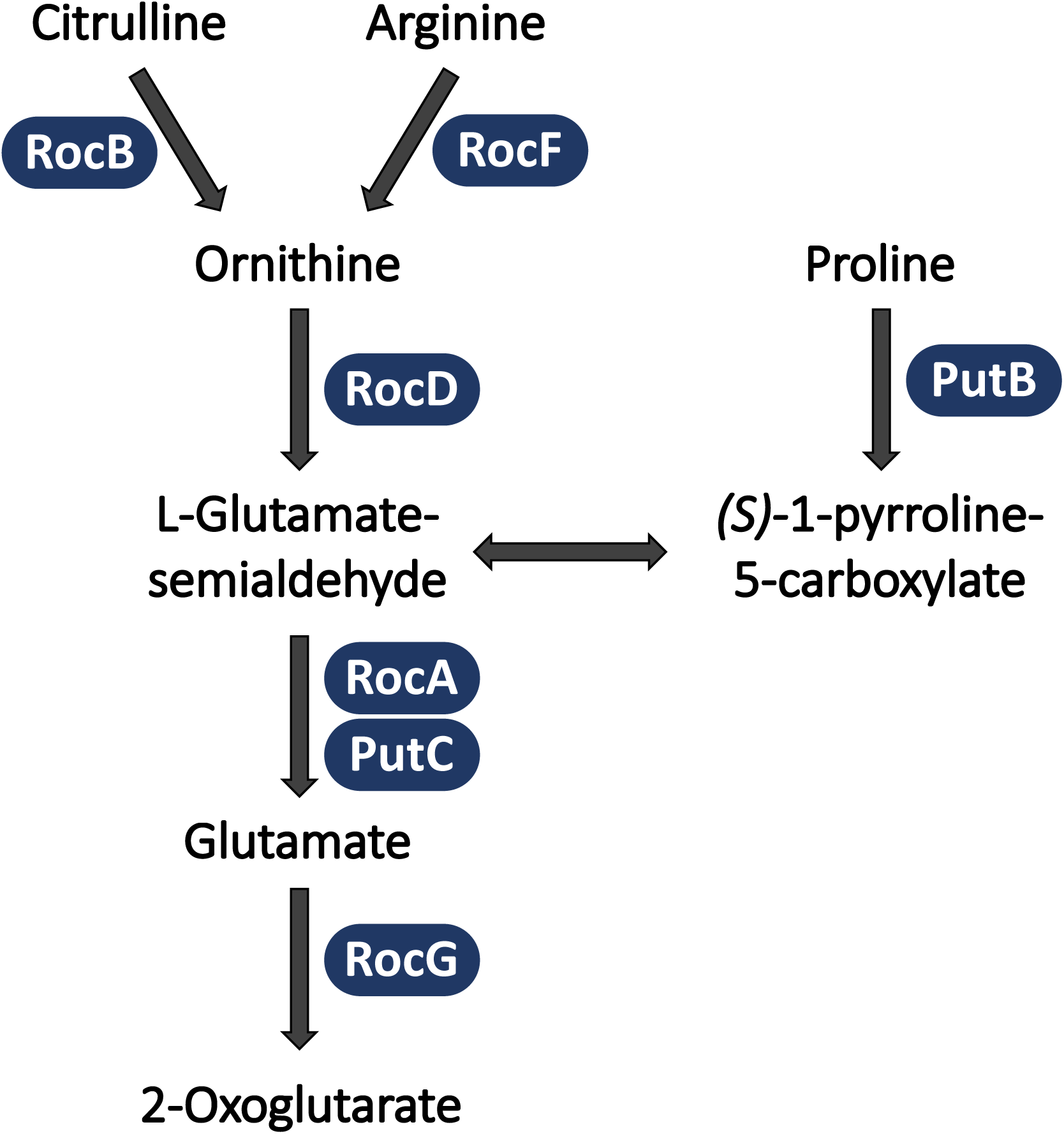
Catabolic routes for citrulline, arginine, and proline in *B. subtilis*. RocB is a N-carbamoyl-L-ornithine hydrolase that catalyzes the first step of citrulline utilization.

The genes for arginine utilization are organized in two gene clusters: The *rocABC* operon encodes an amino acid permease, the enzyme catalyzing the penultimate step in arginine degradation, and an uncharacterized enzyme, RocB (2). This operon is directly clustered with the monocistronic *rocG* gene which encodes the glutamate dehydrogenase which catalyzes the conversion of glutamate to 2-oxoglutarate (3). In addition to RocG, *B. subtilis* encodes a second glutamate dehydrogenase, GudB. This gene is active in non-domesticated strains of *B. subtilis* but inactive due to a duplication of nine base pairs in the laboratory strain 168 (3, 4, 5). The *rocDEF* operon encodes another amino acid permease and the enzymes for the first two steps of arginine utilization (see Fig. 1, 6). The *rocR* gene encoding the transcriptional activator of the genes involved in arginine catabolism is encoded upstream of the *rocDEF* operon, in divergent orientation (2, 6).

Two regulators are involved in the control of arginine degradation, the above-mentioned activator RocR, and the bifunctional activator/ repressor AhrC (2, 6, 7, 8). RocR belongs to a family of transcriptional regulators that activate transcription at a peculiar class of promoters that are recognized by the alternative sigma factor σ^L^. RNA polymerase containing this sigma factor is unable to form the open transcription complex unless transcription is activated by an ATP-hydrolyzing transcription factor (9). RocR is thought to respond to the presence of ornithine and/ or citrulline and to activate expression of the genes of the catabolic pathway in the presence of these intermediates of arginine degradation (10). Both the sigma factor σ^L^ and RocR are essential for the expression of the *roc* genes. The N-terminal domain of RocR acts as an intramolecular repressor domain that controls the activity of the ATP-hydrolyzing central domain, and mutations in this domain as well as in the central domain can result in inducer-independent constitutive activity of RocR 10, 11). AhrC represses the genes for arginine biosynthesis and activates arginine degradation in the presence of arginine (12). The AhrC-repressed arginine biosynthetic pathway is tightly linked to proline biosynthesis and to adaptation of *B. subtilis* to potassium starvation, as both inactivation of the initial genes for proline biosynthesis and severe potassium starvation caused by a deletion of the genes for the low affinity potassium transporters directly select for the inactivation of the *ahrC* gene and the resulting constitutive expression of the arginine biosynthetic pathway (13, 14). Another link between the proline and arginine metabolic pathways is the physical interaction between the (3-hydroxy-)1-pyrroline-5-carboxylate dehydrogenases RocA and PutC (se Fig. 1, 15). Moreover, the two physiologically opposing enzymes RocF (arginase) and ArgF (ornithine carbamoyltransferase) form a counter-enzyme complex to inhibit arginine biosynthesis at high concentrations of the amino acid (16).

Several aspects of arginine degradation and its regulation have so far remained enigmatic. First, the enzyme that catalyzes the first step of citrulline degradation, its conversion to ornithine has not yet been identified. Moreover, the precise mechanism of activation of *roc* gene expression by RocR has not been fully examined, in particular the molecular inducer that triggers RocR activity has not been identified.

We have a long-standing interest in amino acid homeostasis in *B. subtilis* including the metabolism of arginine and its precursor glutamate (11, 13, 17, 18, 19, 20). In this study, we addressed the open questions of the arginine and citrulline catabolic pathway. We demonstrate that the so far uncharacterized enzyme RocB is responsible for the conversion of citrulline to ornithine. Genetic and biochemical data demonstrate that ornithine is the molecular effector that triggers transcription activation by RocR.

## RESULTS

### The initiation of citrulline utilization requires RocB

Proteins encoded by seven genes are thought to be required for the utilization of arginine, ornithine, and citrulline in *B. subtilis*. The corresponding proteins participate in the uptake of these amino acids and their subsequent degradation to 2-oxoglutarate, an intermediate of the citric acid cycle (see Fig. 1). Interestingly, no function has so far been assigned to the RocB protein. Moreover, an enzyme catalyzing the initial step of citrulline utilization has not yet been identified. This brought us to the hypothesis that RocB, a member of the peptidase M20 family, might convert citrulline to ornithine. To test this hypothesis, we tested the growth of a set of *roc* mutants affected in the enzymes of arginine catabolism, for growth in the presence of ammonium or glutamate (as controls), proline, ornithine, citrulline, or arginine as the single source of nitrogen. As expected, all tested strains grew on ammonium or glutamate (see Fig. 2). The wild type strain could also utilize the other tested amino acids. The *rocF* mutant, that lacks arginase, the first enzyme required for arginine degradation, was unable to grow with arginine as the single nitrogen source but grew with proline, ornithine, or citrulline. The arginase reaction yields ornithine which is subsequently converted to glutamate by the ornithine transaminase RocD and the 1-pyrroline-5-carboxylate dehydrogenase RocA. The *rocD* mutant had the “*roc*” phenotype, *i.e*. it could not utilize arginine, ornithine, citrulline as the only nitrogen source. Recently, RocA was found to physically interact with its paralog PutC which is involved in proline utilization (15). Thus, we tested the individual *rocA* and *putC* mutants, GP3727 and GP3749, respectively, as well as the *rocA putC* double mutant GP3398. While the *rocA* and *putC* single mutant strains were able to utilize any of the tested amino acids, the double mutant was unable to grow with proline, ornithine, citrulline or arginine. This observation indicates that the two enzymes that exhibit 69% identity at the amino acid level can take over each other’s function. The glutamate dehydrogenase RocG catalyzes the oxidative deamination of glutamate, the last step of the arginine degradative pathway. The *rocG* mutant was viable with any of the amino acids as the single source of nitrogen since glutamate itself serves as the global amino group donor in the cell. For the *rocB* mutant, we observed growth with all tested amino acids except citrulline. Growth was rescued by expression of plasmid-borne RocB in the mutant strain GP3713 (see Supplemental Figure S1). This suggests that RocB is the missing enzyme for the conversion of citrulline to ornithine.

**Fig. 2.**
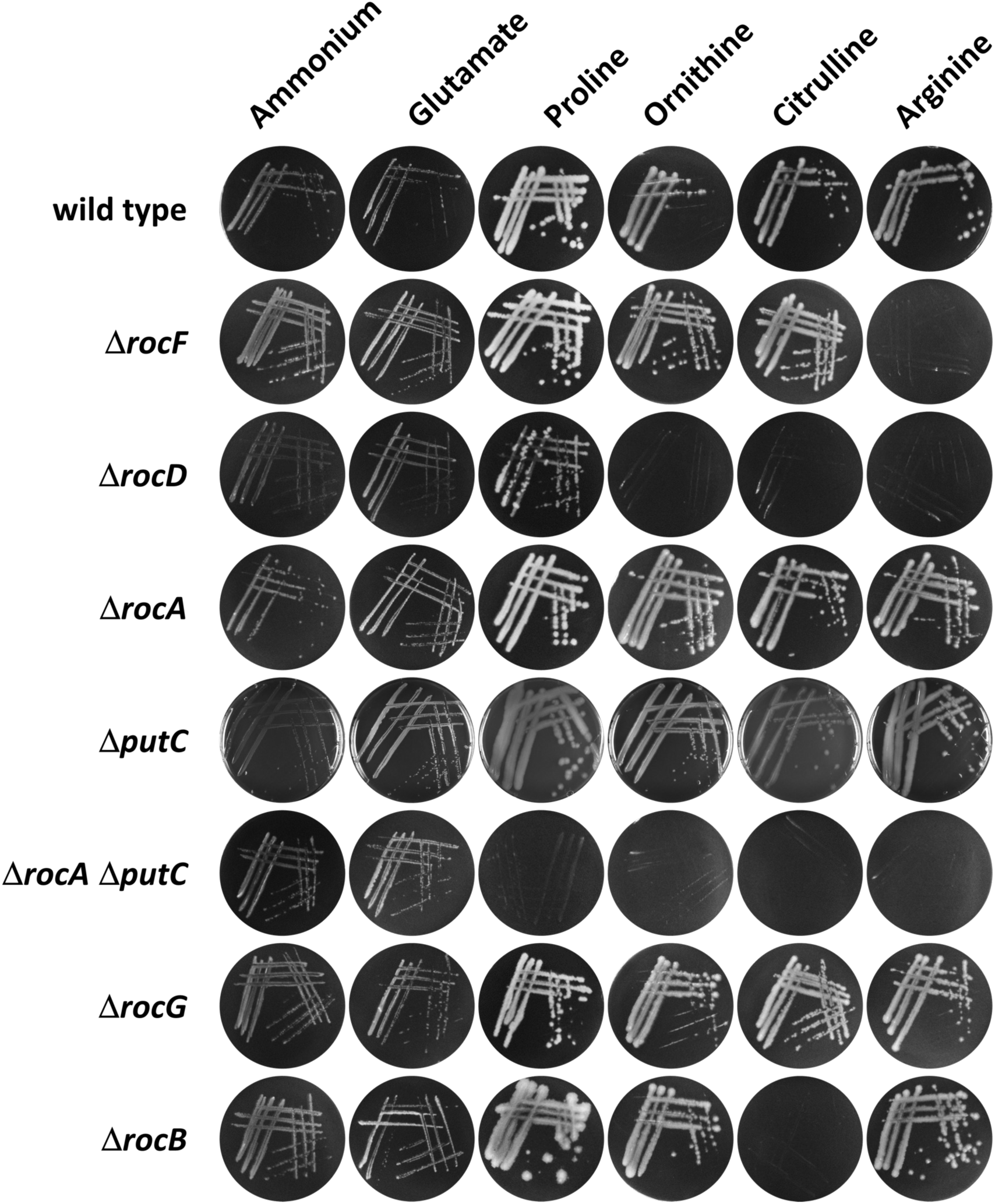
A *rocB* mutant is unable to initiate citrlline utilization. The growth of the wild type strain 168 and of the isogenic mutants deleted for *rocF* (GP655), *rocD* (GP656), *rocA* (GP3747), *putC* (GP3749), *rocA putC* (GP3398), *rocG* (GP726), and *rocB* (GP3720) was tested on Glucose minimal (C Glc) plates with different nitrogen sources. The plates contained 15 mM each of ammoniumsulfate, glutamate, proline, ornithine, citrulline, or arginine. The plates were incubated at 37°C for 48 hours

Taken together, our observations clearly demonstrate that the two highly similar and physically interacting 1-pyrroline-5-carboxylate dehydrogenases can functionally replace each other, and that RocB is specifically required for the introduction of citrulline into the *roc* degradative pathway.

### RocB is a N-carbamoyl-L-ornithine hydrolase

RocB is a member of the M20 peptidase family. For HyuC, a member of this family in *Pseudomonas* sp. NS671, it was shown that the enzyme can release an N-terminal carbamate from N-carbamoyl-L-methionine (21). Since citrulline can be regarded as N-carbamyl-L-ornithine we assumed that the enzyme might release (unstable) carbamate from citrulline. To test the activity of the enzyme, we purified the RocB protein as well as the ornithine transaminase RocD and used both enzymes to determine the formation of pyrroline-5-carboxylate from citrulline in a coupled assay that detects a color change resulting from the formation of a complex of pyrroline-5-carboxylate with 2-aminobenzoate (see Fig. 3A). First, we screened for metal ion requirement by incubating the reaction mix in the presence of magnesium, manganese, or zinc ions. A yellow color indicative of enzymatic activity was only observed in the presence of manganese. Thus, all subsequent assays were performed in the presence of manganese ions. As controls, we performed the assay in the absence of citrulline or with citrulline in the absence of the enzyme (RocB). In both cases, no color change was observed, whereas the solution adopted a yellow color if both citrulline and the enzyme were present in the assay mixture. These results demonstrate that RocB is indeed capable of using citrulline as a substrate for the production of ornithine. To study the enzymatic activity of RocB in more detail, we determined the kinetic parameters of the enzyme. With manganese as divalent metal ion, we observed the formation of pyrroline-5-carboxylate from citrulline (see Fig. 3B). The K_M_ value for citrulline was 7.355 mM l^-1^; the v_max_ at 25°C was 29.78 µmol pyrroline-5-carboxylate (min mmol of enzyme)^-1^. This finding supports the idea that RocB catalyzes the initial step of citrulline utilization.

**Fig. 3.**
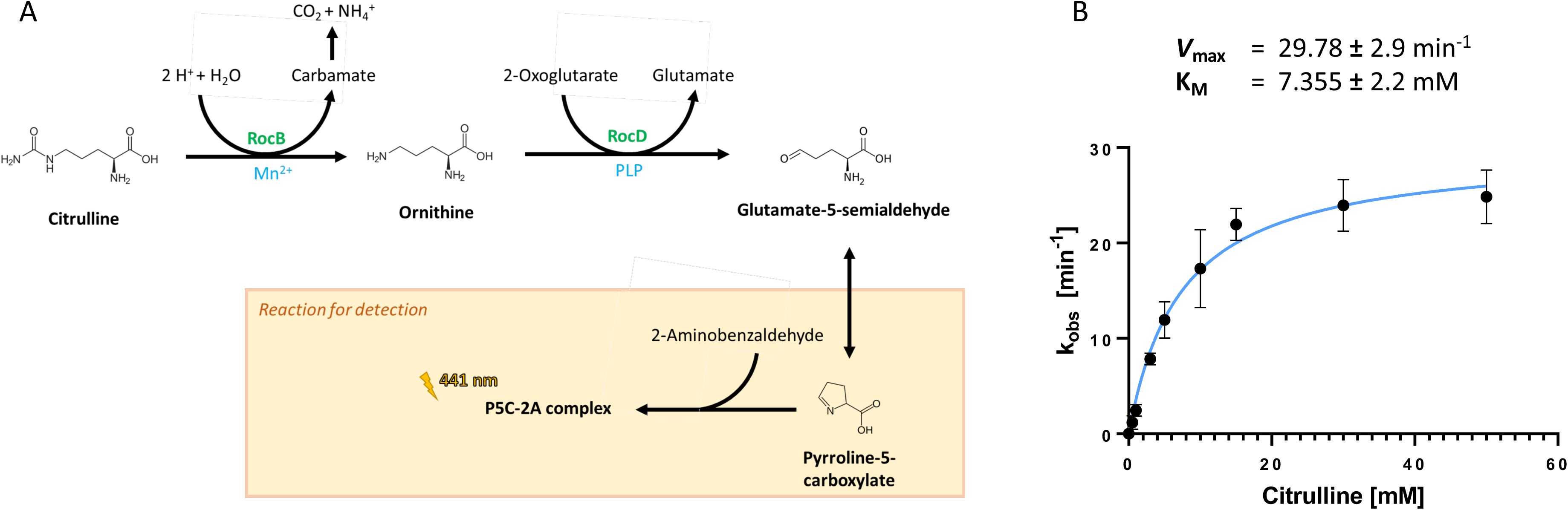
RocB is a N-carbamoyl-L-ornithine hydrolase. **A.** Schematic overview of the coupled RocB-RocD enzyme assay. Citrulline is converted by RocB to ornithine and carbamate, which decays to carbon dioxide and ammonium spontaneously. RocD converts ornithine and 2-oxoglutarate toglutamate and glutamate 5-semialdehyde, which sponteanously converts to pyrroline-5-carboxylate. Together with 2-aminobenzaldehyde, a complex is formed that can be measured at 441 nm. **B**. Kinetic analysis of RocB activity at different citrulline concentrations (0 – 60 mM). The saturation curve was fitted to Michaelis– Menten kinetics using GraphPad Prism (GraphPad Software), and Km and Vmax were calculated.

We conclude that RocB acts as a N-carbamoyl-L-ornithine hydrolase that converts citrulline to ornithine and carbamate.

### Ornithine is the molecular inducer of the *rocDEF* operon by RocR

The genes of the *roc* degradation pathway are subject to regulation by multiple transcription factors. While AhrC is known to directly interact with arginine (12), the molecular inducer activating RocR has remained unknown. Physiologically, all three *roc* amino acids are able to induce the expression of the *rocABC* and *rocDEF* operons via RocR and the alternative sigma factor SigL, and ornithine or citrulline have been proposed to be the molecular effector for RocR (10).

To unequivocally identify the molecular effector of RocR, we made use of a reporter fusion of the *lacZ* gene encoding β-galactosidase to the promoter region of the *rocDEF* operon. The activity of the *rocD* promoter reflects the activity of RocR. We introduced the fusion into our set of isogenic *roc* mutants, and determined the promoter activity after growth in minimal medium containing the *roc* amino acids (see Fig 4). As described previously (10), the promoter was active in the wild type strain GP4039 in the presence of either proline, ornithine, citrulline or arginine, (Fig. 4A). Surprisingly, and in contrast to published data (10), we observed no induction in the presence of proline. In the *rocF* mutant GP4040 that is unable to convert arginine to ornithine, arginine has lost the ability to induce the *rocD* promoter and thus to stimulate RocR activity (Fig. 4B). This suggests that arginine by itself does not act as an effector for RocR. Next, we tested promoter activity in the *rocB* mutant GP4043 that is unable to utilize citrulline. In this strain, both ornithine and arginine induced promoter activity whereas citrulline had no effect (Fig. 4C). This observation is in good agreement with the above conclusion that RocB is required to integrate citrulline in the *roc* degradative pathway. Moreover, the requirement for citrulline conversion to allow activation of the *rocD* promoter indicates that citrulline does not act as effector molecule for RocR. Finally, we also tested the activity of the *rocD* promoter in the *rocB rocF* double mutant GP4044. This mutant lacks the initial steps for the utilization of both arginine and citrulline. As shown in Fig. 4D, only ornithine was able to induce the expression of the *rocD* promoter.

**Fig. 4.**
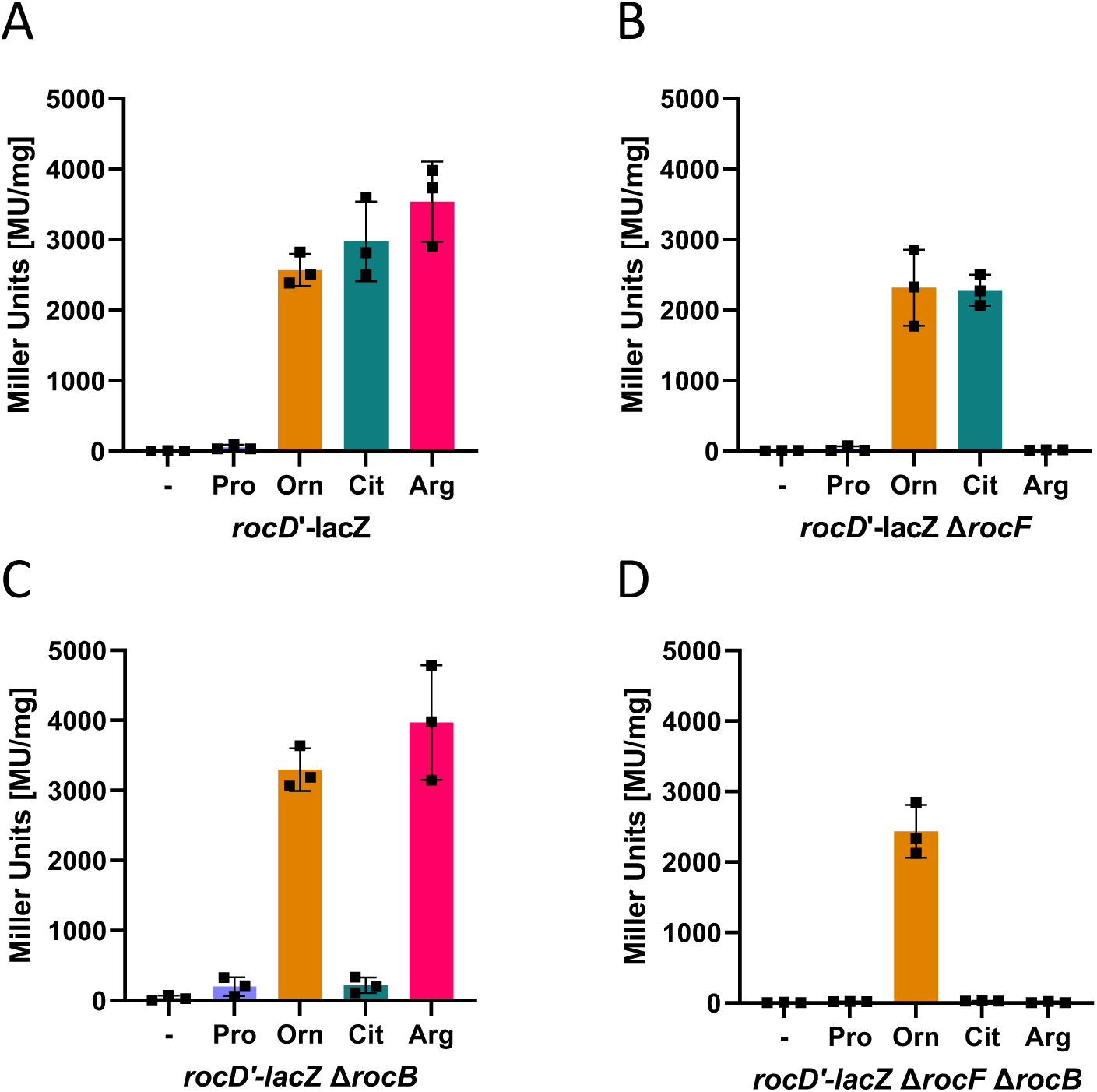
Ornithine triggers transcription activation of the *rocDEF* operon by RocR. To monitor the activation of the *rocD* promoter by RocR, strains that harbour the *rocD-lacZ* reporter gene fusion integrated in the chromosomal *amyE* gene were used. The *rocD-lacZ* reporter gene fusion construct is expressed from the authentic −12/−24 *rocDEF* promoter (Gardan et al., 1995; 1997). Cells were grown in MSSM medium supplemented with 15 mM ammonium sulfate in absence or presence of 15 mM proline, ornithine, citrulline, or arginine. Cultures were grown to early exponential phase (OD_578_ of about 0.6–0.8) and then harvested for β-galactosidase enzyme activity assays. The values for the β-galactosidase activity indicated for each strain represent three independently grown cultures, and for each sample, enzyme activity was determined twice.

Taken together, the results of the promoter assays demonstrate that ornithine is able to induce transcription from the *rocD* promoter whereas neither arginine nor citrulline directly act as inducers. Ornithine and proline share glutamate 5-semialdehyde and glutamate as further intermediates of their catabolic pathways. As proline is unable to induce *rocD* expression, glutamate 5-semialdehyde and glutamate can be excluded as the inducers of *rocD* expression. Thus, ornithine is the molecular inducer and sought-after effector molecule of RocR. The identification of ornithine as the molecular effector of RocR also indicates that RocB directly converts citrulline to ornithine since the inducer can not be formed in the *rocB* mutant in the presence of citrulline.

### Ornithine triggers the ATPase activity of RocR

RocR is a member of the bacterial enhancer binding proteins, which in turn are so-called AAA^+^ ATPases. The enhancer binding proteins are hexameric AAA^+^ ATPases and use their ATPase activities to remodel RNA polymerase complexes containing the alternative sigma factor, σ^54^ (called σ^L^ in *B. subtilis*) to convert the initial closed complex to the open transcription complex (22). The activation of RocR has so far not been studied biochemically. Therefore, we first expressed and purified His-tagged RocR protein. This protein was used in an ATPase activity assay to determine the role of potential effector molecules for RocR activity. The ATPase activity of RocR requires the presence of divalent metal ions. To get more insights into the metal ion requirements of RocR, we perfomed the ATPase assay in the presence of different divalent metal ions. As shown in Fig. 5, manganese (k_obs_ 1.8 min^-1^) was most efficient in stimulating the ATPase activity of RocR, followed by magnesium and cobalt. Since magnesium ions are much more abundant in all cells, we used magnesium for all further assays.

**Fig. 5.**
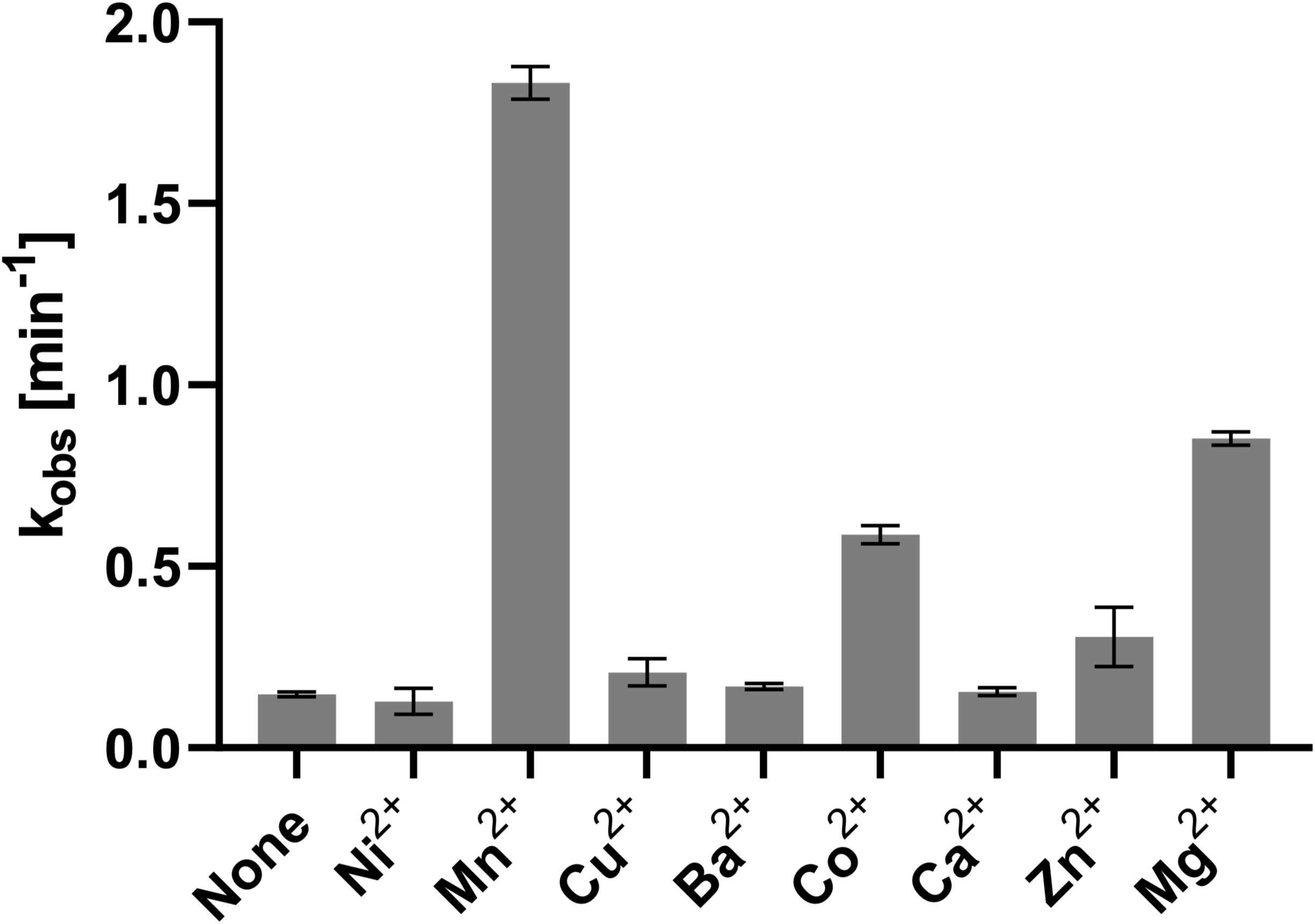
Divalent cations are required for RocR ATPase activity. The activity of RocR was assessed in an *in vitro* ATPase assay. Purified RocR was incubated with ATP in the absence or presence of different divalent cations. The experiment was performed with three independent protein preparations.

Only basal activity (k_obs_ 0.95 min^-1^) was observed in the absence of potential effector molecules (see Fig. 6). Similar results were obtained when the assay was performed in the presence of arginine or citrulline. In contrast, the addition of increasing concentrations of ornithine resulted in an increasing ATPase activity of RocR. This finding is in excellent agreement with the *in vivo* activity assays of the *rocD* promoter that identified ornithine as the molecular effector of RocR (see above).

**Fig. 6.**
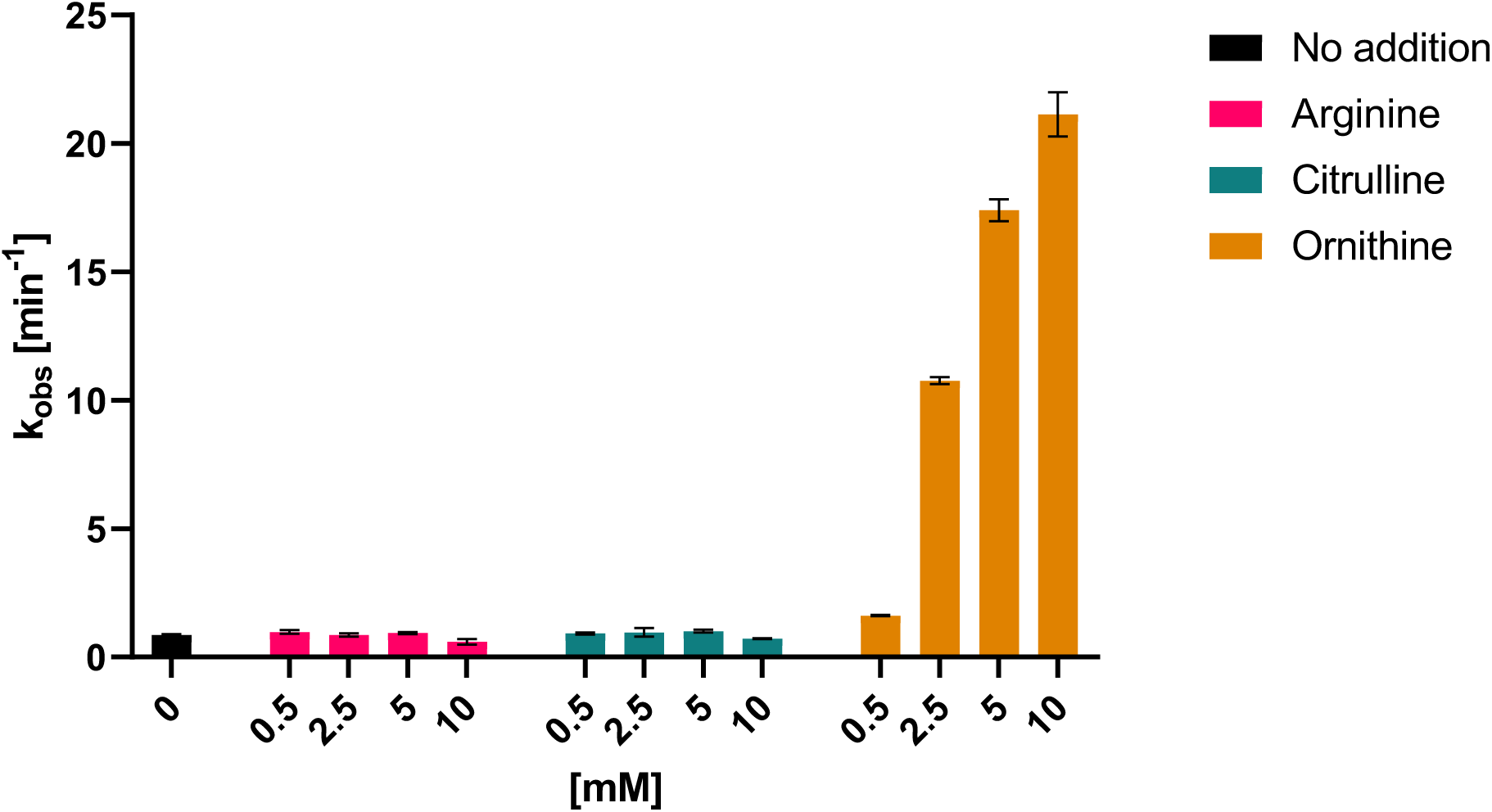
Ornithine triggers the ATPase activity of RocR. The activity of RocR was assessed in an *in vitro* activity assay. Purified RocR was incubated with ATP in the absence or presence of the possible effector molecules arginine, citrulline or ornithine. The experiment was performed with three independent protein preparations.

The RocR protein consists of an N-terminal effector-binding PAS domain, the central domain that binds ATP and interacts with σ^L^, and the C-terminal DNA-binding helix-turn helix domain (10, 23). Several mutations both in the N-terminal and central domains of RocR were found to result in increased transcription of the *roc* genes (10, 11). We were thus interested in the ATPase activity of a truncated RocR protein consisting just of the central domain (amino acids 141 to 400). As described above, the full-length protein exhibited low activity in the absence of an effector and was strongly activated in the presence of ornithine. In contrast, the truncated protein was active irrespective of the presence of potential effector molecules. Even the presence of ornithine did not result in enhanced ATPase activity of the central domain (see Fig. 7). This suggests that the N-terminal domain is indeed mediating the interation with the inducer ornithine, and that it inhibits the activity of the central domain in the absence of the inducer.

**Fig. 7.**
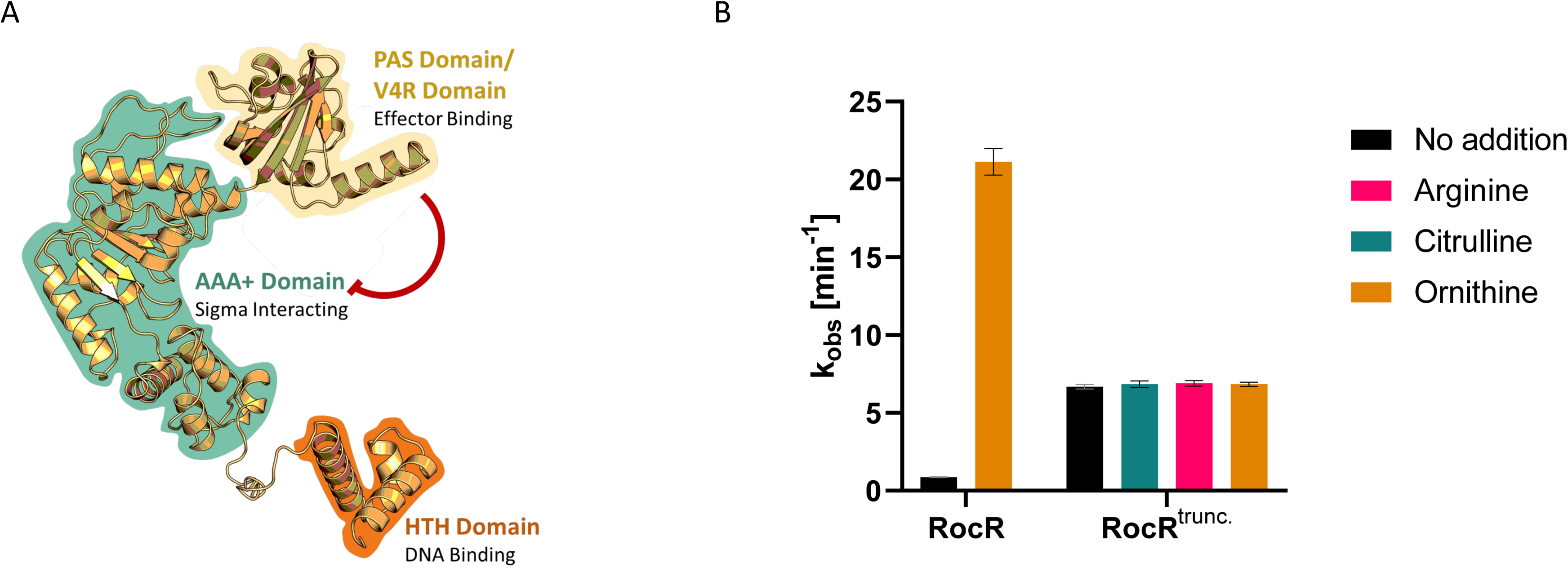
Truncated RocR is active independent of the presence of its inducer. **A.** Structure of the RocR monomer predicted by Alphafold P38022. The protein contains a C-terminal HTH domain (orange) for DNA binding, a central AAA+ ATPase domain (turquoise) for interaction with the sigma factor, and an N-terminal PAS domain/V4R domain (yellow) that regulates the ATPase activity of the protein by effector (ornithine) binding. **B.** The activity of full-length or truncated RocR proteins was assessed in an *in vitro* activity assay. Purified full-length or truncated RocR were incubated with ATP in the absence or presence of the possible effector molecules arginine, citrulline and ornithine. The experiment was performed with three independent protein preparations.

### Binding of an ATP analog affects the oligomeric state of RocR

The bacterial enhancer binding proteins of the AAA^+^ ATPase family are active as hexamers. However, the inactive proteins may also exist in a dimeric state (24). To test the oligomeric state of RocR, we determined the mean particle radius of the protein in the presence and absence of the effector molecule ornithine as well as in the presence of the non-hydrolyzable ATP analog ADP:BeF_3_ by dynamic light-scattering (DLS) (Fig. 8). DLS offers the advantage of studying a macromolecule in its true solution state; chromatography or gel matrices, which can confound interpretations of data by their non-specific interaction with protein, are not present. DLS also indicates the aggregation state of a protein under a given set of experimental conditions. In the absence of any low-molecular weight effector, Apo-RocR had an estimated molecular weight of 121 kDa. Based on the molecular weight of the His_6_-RocR monomer of 55 kDa, this likely corresponds to a dimer. The addition of ornithine resulted in an estimated molecular weight of 123 kDa, indicating that the presence of ornithine did not affect the oligomeric state of the protein. The addition of the ATP analog ADP:BeF_3_ resulted in the formation of high molecular weight assemblies of RocR. The putative donut-like structure of the active AAA^+^ ATPases does not allow to derive a precise molecular weight from the observed particle radius. We assume that the large assemblies correspond to the active hexamers that have been observed for other AAA^+^ ATPases (24). Similar-sized large assemblies were observed if RocR was incubated with both ADP:BeF_3_ and ornithine suggesting that ornithine does not affect the oligomeric state of RocR but rather the activity of the ATP-bound functional hexamer.

**Fig. 8.**
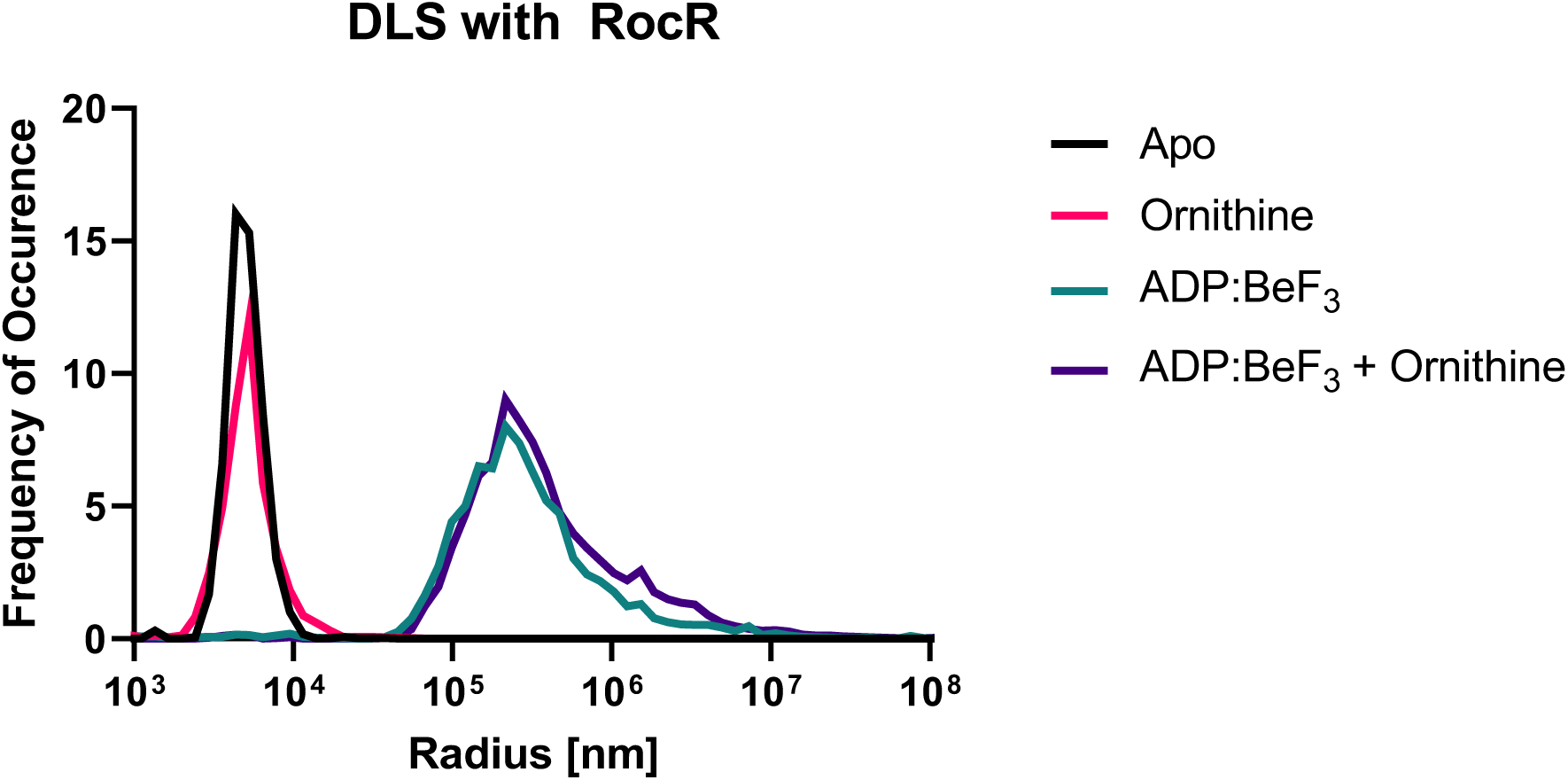
The oligomerization state of RocR is altered in the presence of an ATP-analogue. For the investigation of the oligomerization state of RocR, the mean particle size of the purified protein was analyzed by DLS. As a control, RocR was analyzed in its apo state (black).

## DISCUSSION

The results presented in this study demonstrate the central role of ornithine in the utilization of arginine and citrulline in *B. subtilis*. While all three amino acids can feed into the *roc* catabolic pathway, arginine and citrulline are first converted to ornithine. This metabolite is the central intermediate of the pathway and acts also as the molecular inducer that triggers the activity of the transcription factor RocR (see Fig. 9).

**Fig. 9.**
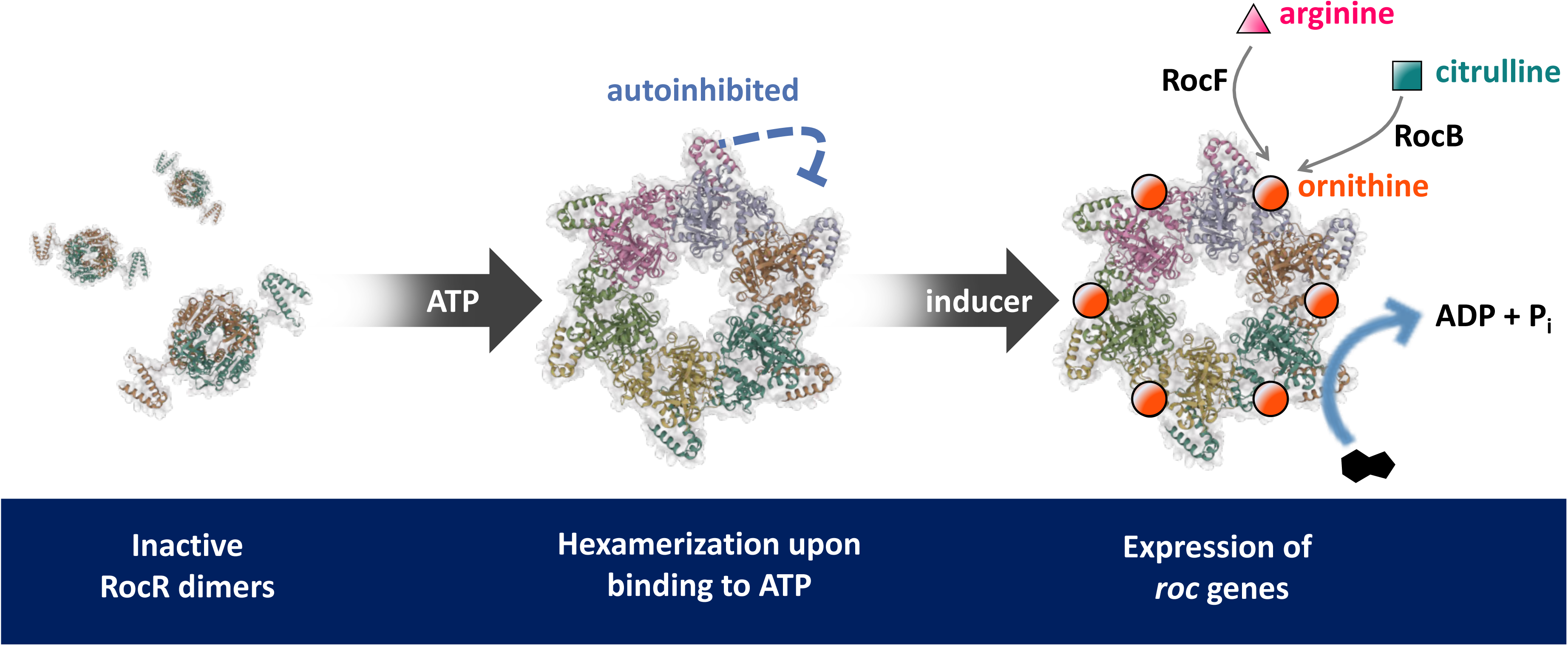
The molecular events that lead to the activation of RocR. In its apo form, RocR exists as a dimer. In the presence of ATP it forms hexamers, but the enzyme complex is autoinhibited in the absence of an inducer. Only in the presence of ornithine RocR exhibits ATPase activity, inducing *roc* gene expression.

The investigation of mutant phenotypes with respect to amino acid utilization as nitrogen sources yielded two important insights. First, the paralogous and physically interacting 1-pyrroline-5-carboxylate dehydrogenases RocA and PutC can functionally replace each other in the utilization of proline and arginine. This suggests that both enzymes are present in sufficient amounts in cells growing in minimal medium even in the absence of the cognate amino acid that could serve as inducer. Interactions between paralogous proteins have been observed in other cases such as the RNases J1 and J2 or the UDP-N-acetylglucosamine 1-carboxyvinyltransferases MurAA and MurAB, even though the enzymes are active independently from each other in the absence of the interaction partner (15, 25). Second, RocB is a N-carbamoyl-L-ornithine hydrolase that is responsible for the so far non-annotated conversion of citrulline to ornithine. This reaction fills the last gap in our knowledge of the *roc* degradative pathway. The ability to use citrulline as a nutrient seems to be limited to a rather small number of members of the Firmicutes as judged from the phylogenetic distribution of the RocB protein, and RocB is the first functionally characterized member of COG4187 (26). Both *roc* operons encode highly similar amino acid permeases. It is tempting to speculate that RocC is more important for the uptake of citrulline whereas RocE might be more relevant for arginine and ornithine transport. However, given the presence of and additional ABC transporter for arginine uptake and the low specificity of amino acid permeases that often exhibit significant substrate promiscuity and transport multiple amino acids (19, 20), this issue must remain open in the context of this study.

Our study also reveals how the transcription factor RocR is activated in a two-step mechanism. First, the binding of ATP triggers the conversion of the inactive dimer to the hexamer. However, in the hexamer the N-terminal domain of RocR inhibits the activity of the central ATP-hydrolyzing domain of the protein (10, 11), and ATP hydrolysis and transcription activation can only occur upon binding of the effector molecule ornithine as shown by *in vitro* ATPase and *in vivo rocD* promoter activity assays (see Fig. 9). In good agreement with this autoinhibition by the N-terminal domain is also the observed ATPase activity of the isolated central domain of RocR irrespective of the presence of ornithine. It is interesting to note that the limitation of protein activity by an autoinhibitory domain is not unprecedented. The *B. subtilis* diadenylate cyclase CdaS can form enzymatically inactive hexamers. Only changes in the N-terminal domain of the protein sterically allow the formation of highly active dimers (27).

Two transcription factors are involved in the regulation of arginine metabolism. The bifunctional AhrC protein acts as repressor for the biosynthetic genes and as activator for the degradative genes (7, 8). AhrC responds to the presence of arginine which acts as a co-repressor and co-activator by allowing binding of AhrC to its target sites in the upstream regions of arginine metabolic genes and operons. In contrast, the RocR activator which is required for the transcription of the degradative *roc* genes, uses ornithine as its cofactor that triggers ATP hydrolysis and productive interaction with the SigL-containing RNA polymerase. Thus, AhrC uses arginine as the single specific effector whereas ornithine is a more general effector molecule for RocR activity as it is the first common intermediate of arginine and citrulline degradation. Thus, the use of RocR in addition to AhrC allows a response not only to arginine bit also to the non-proteinogenic amino acids citrulline and ornithine which are also degraded by the enzymes encoded by the *roc* genes. Clearly, RocR is required for transcription activation using the RNA polymerase containing the alternative sigma factor SigL. However, AhrC has a completely different structure and is not likely to functionally interact with this form of RNA polymerase. It is therefore tempting to speculate that AhrC activates transcription at a second, SigA-dependent promoter just in response to arginine availability. This idea is supported by the overlap of the SigL-dependent promoter and the AhrC binding site upstream of the *rocABC* operon (8) which is not compatible with AhrC-mediated transcription activation at this promoter. In this way, the two promoters could allow appropriate regulation of the *roc* genes in the presence of arginine alone or its non-proteinogenic relatives.

## MATERIALS AND METHODS

### Strains, media and growth conditions

*E. coli* DH5α (Sambrook et al., 1989) was used for cloning. All *B. subtilis* strains used in this study are derivatives of the laboratory strain 168. They are listed in Table 1. *B. subtilis* and *E. coli* were grown in Luria-Bertani (LB) or in sporulation (SP) medium (28, 29). For growth assays, *B. subtilis* was cultivated in MSSM medium (30). MSSM is a modified SM medium in which KH_2_PO_4_ was replaced by NaH_2_PO_4_ and KCl was added as indicated (30). The media were supplemented with ampicillin (100 µg/ml), kanamycin (10 µg/ml), chloramphenicol (5 µg/ml), spectinomycin (150 µg/ml), tetracycline (12.5 µg/ml) or erythromycin and lincomycin (2 and 25 µg/ml, respectively) if required.

### DNA manipulation and transformation

All commercially available restriction enzymes, T4 DNA ligase and DNA polymerases were used as recommended by the manufacturers. DNA fragments were purified using the QIAquick PCR Purification Kit (Qiagen, Hilden, Germany). DNA sequences were determined by the dideoxy chain termination method (28). Standard procedures were used to transform *E. coli* (28), and transformants were selected on LB plates containing ampicillin (100 µg/ml). *B. subtilis* was transformed with plasmid or chromosomal DNA according to the two-step protocol described previously (Kunst et al., 1995). Transformants were selected on SP plates containing chloramphenicol (Cm 5 µg/ml), kanamycin (Km 10 µg/ml), spectinomycin (Spc 150 µg/ml), tetracycline (Tet 12,5 µg/ml) or erythromycin plus lincomycin (Em 2 µg/ml and Lin 25 µg/ml).

**Table 1.** *B. subtilis* strains used in this study

| Strain | Genotype | Source or Reference |
| --- | --- | --- |
| 168 | *trpC2* | Laboratory collection |
| GP655 | *trpC2 ΔrocF::aphA3* | QB5585 → 168 |
| GP656 | *trpC2 ΔrocD::aphA3* | QB5618 → 168 |
| GP726 | *trpC2 rocG::aphA3* | This study |
| GP3398 | *trpC2 ΔputC::aphA3 ΔrocA::tet* | GP3727 → GP3749 |
| GP3720 | *trpC2 ΔrocB::cat* | This study |
| GP3722 | *trpC2 ΔrocB::tet* | This study |
| GP3727 | *trpC2 ΔrocA::tet* | This study |
| GP3747 | *trpC2 ΔrocA::aphA3* | This study |
| GP3749 | *trpC2 ΔputC::aphA3* | This study |
| GP4039 | *trpC2 amyE::rocD'-lacZ-cat* | QB5556 → 168 |
| GP4040 | *trpC2 amyE::rocD'-lacZ-cat ΔrocF::aphA3* | QB5556 → GP655 |
| GP4043 | *trpC2 amyE::rocD'-lacZ-cat ΔrocB::tet* | GP3722 → GP4039 |
| GP4044 | *trpC2 amyE::rocD'-lacZ-cat ΔrocF::aphA3*<br>*ΔrocB::tet* | GP3722 → GP4040 |
| QB5556 | *trpC2 amyE ::(rocD-lacZ cat)* | 6 |
| QB5585 | *trpC2 rocF::aphA3 amyE::rocD'-lacZ cat* | 10 |
| QB5618 | *trpC2 rocD::aphA3 amyE::rocD'-lacZ cat* | 10 |

### Construction of mutant strains by allelic replacement

Deletion of the *putC*, *rocA*, *rocB*, and *rocG* genes was achieved by transformation of *B. subtilis* 168 with a PCR product constructed using oligonucleotides to amplify DNA fragments flanking the gene of interest and intervening resistance cassette specifying antibiotiresistance to tetracycline as described previously (31). The integrity of the regions flanking the integrated resistance cassette was verified by sequencing PCR products of about 1,100 bp amplified from chromosomal DNA of the resulting mutant strains (see Table 1).

### Phenotypic analysis

Quantitative studies of *lacZ* expression in *B. subtilis* were performed as follows: cells were grown in MSSM medium supplemented with KCl at different concentrations as indicated. Cells were harvested at OD_600_ of 0.5 to 0.8. β-Galactosidase specific activities were determined with cell extracts obtained by lysozyme treatment as described previously (29). One unit of β-galactosidase is defined as the amount of enzyme which produces 1 nmol of o-nitrophenol per min at 28°C.

### Plasmid constructions

To allow overexpression of the *rocB* gene in *B. subtilis*, we constructed plasmids pGP3731. The *rocB* gene was amplified using oligonucleotides that added BamHI and PstI sites to the ends of the fragment and cloned into the expression vector pBQ200 (32) linearized with the same enzymes.

For the purification of RocB and RocD, each carrying a Strep-tag for affinity purification, we amplified the *rocB* and *rocD* alleles using chromosomal DNA of *B. subtilis* 168 as the template and appropriate oligonucleotides that attached specific restriction sites to the fragment. The enzymes KpnI and BamHI were used for integration into the vector pGP172 (33) that encodes an N-terminal Strep-tag. The resulting plasmids were pGP3794 for *rocB* and pGP3795 for *rocD*.

For purification of RocR, plasmid pGP3855 for inducible expression of RocR carrying a His-tag at its N-terminus was constructed as follows. The *rocR* gene was amplified using primers that add BamHI and PstI sites to the ends of the fragment and cloned into the expression vector pWH844 (34) linearized with the same enzymes. To purify the central domain of RocR which bind ATP and is responsible for the interaction with σ^L^, we cloned the region of the *rocR* gene that corresponds to amino acids 141 to 400 into pET-SUMO (Invitrogen, Germany). For this purpose, the *rocR* gene fragment was amplified using primers that add BamHI and NotI sites to the ends of the fragment and cloned into the vector linearized with the same enzymes. The resulting plasmid was pGP3721.

### Protein expression and purification

For the purification of Strep-tagged RocB and RocD, *E. coli* BL21(DE3) was transformed with plasmids pGP3794 and pGP3795, respectively. Expression of the recombinant proteins was induced by the addition of isopropyl 1-thio-β-D-galactopyranoside (final concentration, 1 mM) to exponentially growing cultures (OD_600_ of 0.8) of *E. coli* carrying the relevant plasmid. Strep-tagged proteins were purified in Buffer W (100 mM Tris-HCl, 150 mM NaCl, 1 mM Na_2_EDTA, pH 8.0). Cells were lysed by four passes at 18,000 p.s.i. through an HTU DIGI-F press (G. Heinemann, Germany). After lysis, the crude extracts were centrifuged at 100,000 × *g* for 60 min and then passed over a StrepTactin column (IBA, Göttingen, Germany) for affinity binding of Strep-tagged proteins. The protein was eluted with D-desthiobiotin (2.5 mM). After elution, the fractions were tested for the desired protein using SDS-PAGE. The protein samples were stored at−80 °C until further use (but no longer than 3 days).

For the purification of 6xHis-RocR and 6xHis-SUMO-RocR^trunc^, *E. coli* BL21(DE3) was transformed with plasmids pGP3855 or pGP3721, respectively. Briefly, cells were grown in 2YT medium and expression was induced by the addition of isopropyl 1-thio-β-D-galactopyranoside (final concentration, 0.5 mM) to exponentially growing cultures (OD_600_ of 0.7). The cultures were incubated at 16°C and 210 rpm overnight. Cells were disrupted by 7 passes at 15,000 p.s.i through a microfluidizer (Microfluids, Jena) in 50 mM Tris (pH7.5), 500 mM NaCl and 20 mM imidazole. The crude extract was centrifuged at 35,000 g for 30 minutes and then passed over Ni Sepharose HP (GE Healthcare, Solingen). The target proteins were eluted with an imidazole gradient. The fractions were tested for the desired protein using 17.5% SDS-PAGE. The fractions of interest were pooled and supplemented with EDTA (final concentration, 10 mM). The truncated construct was furthermore treated with 1:100 (w/w) SUMO protease. The resulting protein pools for both proteins were dialyzed in 10 mM Tris (pH7.5), 200 mM NaCl overnight. To remove the cleaved-off tag from the truncated RocR construct, a second Ni Sepharose purification step was employed.

The protein concentration was determined using the Bio-Rad dye binding assay and bovine serum albumin as the standard (35).

### RocB enzyme assay

The RocB activity was measured using a coupled assay of pyrroline-5-carboxylate (P5C) formation (see Fig. 3A, adapted from (36)). Formation of the P5C-*o*-aminobenzaldehyde complex was measured by the absorption at 441 nm, which was recorded over time with an Epoch 2 Microplate Spectrophotometer (BioTek Instruments). All reactions were performed in triplicates of 60 µl each at 25°C in PBS, 2 mM MnCl_2_, 80 µM PLP, 10 mM 2-oxoglutarate, 5 mM *o*-aminobenzaldehyde in 20% ethanol. RocD and RocB were added at concentrations of 1 µM each. The formation of the P5C-*o*-aminobenzaldehyde complex (µmol per minute (k_obs_)) was calculated using

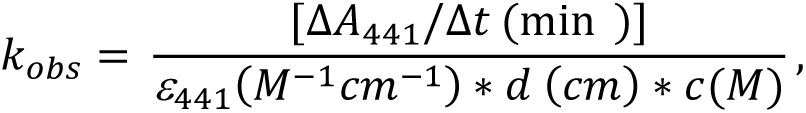

where ΔA_441_/Δt is the slope of P5C-*o*-aminobenzaldehyde complex formation, ε_441_ is the extinction coefficient of the P5C-*o*-aminobenzaldehyde complex, d is the optical pathlength and c is the protein concentration. The millimolar extinction coefficient is 2.71 (37).

### ATPase assay

The ATPase activity of the full-length and truncated RocR proteins was monitored employing a standard enzymatic assay following NADH absorption at 340 nm (38). The decrease of the NADH absorption was recorded with a VICTOR Nico Multimode Microplate Reader (Perkin Elmer, Rodgau, Germany). All reactions were performed in triplicates of 150 µl at room temperature. Each reaction was performed in 25 mM Tris/HCl (pH 7.5), 150 mM NaCl, 15 mM of MgCl_2_ or other metal ions as indicated, 250 nM NADH, 500 nM phosphoenolpyruvate, 6-8.3 U/ml pyruvate kinase, 9-14 U/ml lactate dehydrogenase and 15 mM ATP. To obtain suitable reaction velocities, RocR was used at a concentration of 2 µM (metal ion specificity test), 0.5 µM (stimulation test), or 1 µM (truncated RocR). Arginine, citrulline and ornithine were added as indicated. The ATP consumption per minute (k_obs_) was calculated as described (38).

### Dynamic light scattering

Dynamic light-scattering studies on RocR were performed using a SpectroLight 600/610 (Xtal Concepts GmbH, Hamburg). RocR samples at a concentration of 0.5 mg ml^-1^ were prepared in solutions of 25 mM Tris (pH 7.5), 150 mM NaCl, 15 mM MgCl_2_. Ornithine and ADP:BeF_3_ were added to final concentrations of 10 mM as indicated. Prior to the experiment, all samples were centrifuged at 16.873 x g at 10°C for 30 minutes to eliminate any large aggregates. The reported values are averages of 20 scans of 20 seconds each.

## ACKNOWLEDGEMENTS

This work was supported by a grant of the Deutsche Forschungsgemeinschaft (DFG) within the Priority Program SPP1879 (to R.F. and J.S.). The funders had no role in study design, data collection, analysis and interpretation, decision to submit the work for publication, or preparation of the manuscript.

## Author contributions

Design of the study: R.W. and J.S. Experimental work: R.W., G.A., C.H. and T.G. Data analysis: R.W., T.G., R.F., and J.S.. Wrote the paper: R.W. and J.S.

## REFERENCES

1. Débarbouillé M, Martin-Verstraete I, Kunst F, Rapoport G. 1991. The *Bacillus subtilis sigL* gene encodes an equivalent of sigma 54 from gram-negative bacteria. Proc Natl Acad Sci U S A 88:9092–9096.

2. Calogero S, Gardan R, Glaser P, Schweizer J, Rapoport G., Débarbouillé M. 1994. RocR, a novel regulatory protein controlling arginine utilization in *Bacillus subtilis*, belongs to the NtrC/NifA family of transcriptional activators. J Bacteriol 176:1234–1241.

3. Belitsky BR, Sonenshein AL. 1998. Role and regulation of *Bacillus subtilis* glutamate dehydrogenase genes. J Bacteriol 180:6298–6305.

4. Zeigler DR, Prágai Z, Rodriguez S, Chevreux B, Muffler A, Albert T, Bai B, Wyss M, Perkins JB. 2008. The origins of 168, W23, and other *Bacillus subtilis* legacy strains. J Bacteriol 190:6983–6995.

5. Gunka K, Tholen S, Gerwig J, Herzberg C, Stülke J, Commichau FM. 2012. A high-frequency mutation in *Bacillus subtilis*: requirements for the decryptification of the *gudB* glutamate dehydrogenase gene. J Bacteriol 194:1036–1044.

6. Gardan R, Rapoport G, Débarbouillé M. 1995. Expression of the *rocDEF* operon involved in arginine catabolism in *Bacillus subtilis*. J Mol Biol 249:843–856.

7. Klingel U, Miller CM, North AK, Stockley PG, Baumberg S. 1995. A binding site for activation by the *Bacillus subtilis* AhrC protein, a repressor/ activator of arginine metabolism. Mol Gen Genet 248:329–340.

8. Miller CM, Baumberg S, Stockley PG. 1997. Operator interactions by the *Bacillus subtilis* arginine repressor/ activator, AhrC: novel positioning and DNA-mediated assembly of a transcriptional activator at catabolic sites. Mol Microbiol 26:37–48.

9. Shingler V. 2011. Signal sensory systems that impact σ^54^-dependent transcription. FEMS Microbiol Rev 35:425–436.

10. Gardan R, Rapoport G, Débarbouillé M. 1997. Role of the transcriptional activator RocR in the arginine-degradation pathway of *Bacillus subtilis*. Mol Microbiol 24:825–837.

11. Zaprasis A, Hoffmann T, Wünsche G, Flórez LA, Stülke J, Bremer E. 2014. Mutational activation of the RocR activator and of a cryptic *rocDEF* promoter bypass loss of the initial steps of proline biosynthesis in *Bacillus subtilis*. Environ Microbiol 16:701–717.

12. Garnett JA, Baumberg S, Stockley PG, Phillips SEV. 2007. Structure of the C-terminal effector binding domain of AhrC bound to its corepressor arginine. Acta Crystallogr Sect F Struct Biol Cryst Commun 63:918–921.

13. Gundlach J, Herzberg C, Hertel D, Thürmer A, Daniel R, Link H, Stülke J. 2017. Adaptation of *Bacillus subtilis* to life at extreme potassium limitation. mBio 8:e00861–17.

14. Stecker D, Hoffmann T, Link H, Commichau FM, Bremer E. 2022. L-proline synthesis mutants of *Bacillus subtilis* overcome osmotic sensitivity by genetically adapting L-arginine metabolism. Front Microbiol 16:908304.

15. O’Reilly FJ, Graziadei A, Forbrig C, Bremenkamp R, Charles K, Lenz S, Elfmann C, Fischer L, Stülke J, Rappsilber J. 2023. Protein complexes in cells by AI-assisted structural proteomics. Mol Syst Biol 19:e11544.

16. Isally IM, Isally AS. 1074. Control of ornithine carbamoyltransferase activity by arginase in *Bacillus subtilis*. Eur J Biochem 49:485–495.

17. Ludwig H, Meinken C, Matin A, Stülke J. 2002. Insufficient expression of the *ilv-leu* operon encoding enzymes of branched-chain amino acids biosynthesis limits growth of a *Bacillus subtilis ccpA* mutant. J Bacteriol 184:5174–5178.

18. Commichau FM, Gunka K, Landmann JJ, Stülke J. 2008. Glutamate metabolism in *Bacillus subtilis*: gene expression and enzyme activities evolved to avoid futile cycles and to allow rapid responses to perturbations of the system. J Bacteriol 190:3557–3564.

19. Klewing A, Koo BM, Krüger L, Poehlein A, Reuß D, Daniel R, Gross CA, Stülke J. 2020. Resistance to serine in *Bacillus subtilis*: identification of the serine transporter YbeC and of a metabolic network that links serine and threonine metabolism. Environ Microbiol. 22:3937–3949.

20. Krüger L, Herzberg C, Rath H, Pedreira P, Ischebeck T, Poehlein A, Gundlach J, Daniel R, Völker U, Mäder U, Stülke J. 2021. Essentiality of c-di-AMP in *Bacillus subtilis*: bypassing mutations converge in potassium and glutamate homeostasis. PLoS Genet 17:e1009092.

21. Watabe K, Ishikawa T, Mukohara Y, Nakamura H. 1992. Cloning and sequencing of the genes involved in the conversion of 5-substituted hydantoins to the corresponding L-amino acids from the native plasmid of *Pseudomonas* sp. Strain NS671. J Bacteriol 174:962–969.

22. Gao F, Danson AE, Ye F, Jovanovich M, Buck M, Zhang X. 2020. Bacterial enhancer binding proteins – AAA+ proteins in transcription activation. Biomolecules 10:351.

23. Studholme DJ, Dixon R. 2003. Domain architectures of sigma54-dependent transcriptional activators. J Bacteriol 185:1757–1767.

24. Bush M, Dixon R. 2012. The role of bacterial enhancer binding proteins as specialized activators of σ^54^-dependent transcription. Microbiol Mol Biol Rev 76:497–529.

25. Mathy N, Hébert A, Mervelet P, Bénard L, Dorléans A, de la Sierray-Gallay IL, Noirot P, Putzer H, Condon C. 2010. *Bacillus subtilis* ribonucleases J1 and J2 form a complex with altered enzyme behaviour. Mol Microbiol 75:489–498.

26. Galperin MY, Wolf YI, Makarova KS, Vera Alvarez R, Landsman D, Koonin EV. 2021. COG database update: focus on microbial diversity, model organisms, and widespread pathogens. Nucleic Acids Res 49:D274–D281.

27. Mehne FM, Schröder-Tittmann K, Eijlander RT, Herzberg C, Hewitt L, Kaever V, Lewis RJ, Kuipers OP, Tittmann K, Stülke J. 2014. Control of the diadenylate cyclase CdaS in *Bacillus subtilis*: an autoinhibitory domain limits cyclic di-AMP production. J Biol Chem 289:21098–21107.

28. Sambrook J, Fritsch EF, Maniatis T. 1989. Molecular cloning: a laboratory manual, 2nd ed. Cold Spring Harbor Laboratory, Cold Spring Harbor, N.Y.

29. Kunst F, Rapoport G. 1995. Salt stress is an environmental signal affecting degradative enzyme synthesis in *Bacillus subtilis*. J Bacteriol 177:2403–2407.

30. Gundlach J, Krüger L, Herzberg C, Turdiev A, Poehlein A, Tascon I, Weiss M, Hertel D, Daniel R, Hänelt I, Lee VT, Stülke J. 2019. Sustained sensing in potassium homeostasis: cyclic di-AMP controls potassium uptake by KimA at the levels of expression and activity. J Biol Chem 294:9605–9614.

31. Diethmaier C, Newman JA, Kovács AT, Kaever V, Herzberg C, Rodrigues C, Boonstra M, Kuipers OP, Lewis RJ, Stülke J. 2014. The YmdB phosphodiesterase is a global regulator of late adaptive responses in *Bacillus subtilis*. J Bacteriol 196:265–275.

32. Martin-Verstraete I, Débarbouillé M, Klier A, Rapoport G. 1994. Interactions of wild-type and truncated LevR of *Bacillus subtilis* with the upstream activating sequence of the levanase operon. J Mol Biol 241:178–192.

33. Merzbacher M, Detsch C, Hillen W, Stülke J. 2004. *Mycoplasma pneumoniae* HPr kinase/ phosphorylase: assigning functional roles to the P-loop and the HPrK/P signature sequence motif. Eur J Biochem 271:367–374.

34. Schirmer F, Ehrt S, Hillen W. 1997. Expression, inducer spectrum, domain structure, and function of MopR, the regulator of phenol degradation in Acinetobacter calcoaceticus NCIB8250. J Bacteriol 179:1329–1336.

35. Bradford MM. 1976. A rapid and sensitive method for the quantification of microgram quantities of protein utilizing the principle of protein-dye binding. Anal Biochem 72:248–254.

36. Dendinger S, Brill WJ. 1970. Regulation of proline degradation in *Salmonella typhimurium*. J Bacteriol 103:144–152.

37. Strecker HJ. 1965. Purification and properties of rat liver ornithine delta-transaminase. J Biol Chem 240:1225–1230.

38. Agarwal KC, Miech RP, Parks RE. 1978. Guanylate kinases from human erythrocytes, hog brain, and rat liver. Meth Enzymol 51: 483–490.

