## Supplemental Figure S1 for "Ornithine is the central intermediate in the arginine degradative pathway and its regulation in *Bacillus subtilis*"

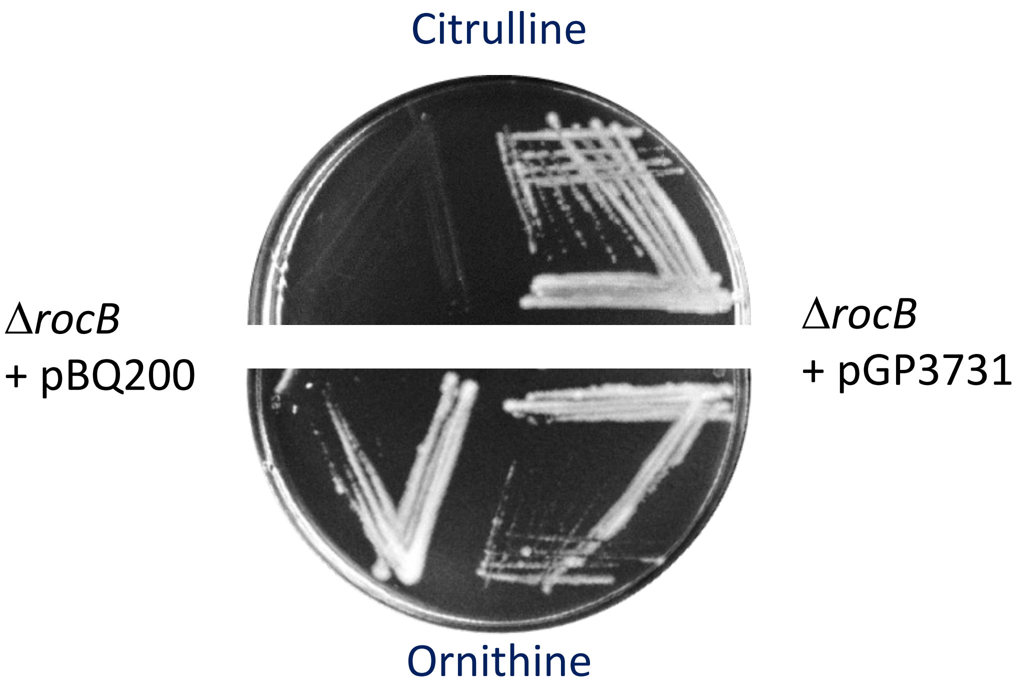


**Plasmid-borne RocB restores citrulline utilization of a *rocB* mutant.** The growth of the *rocB* deletion mutant GP3720 carrying either the empty vector pBQ200 or pGP3731 for constitutive expression of *rocB* on glucose minimal (C Glc) plates with 15 mM citrulline or ornithine as nitrogen source was tested. The plates were incubated at 37°C for 48 hours.

Supplemental Figure S1
